# SIRT1 mediates the antagonism of Wnt/β-catenin pathway by vitamin D in colon carcinoma cells

**DOI:** 10.1101/2024.01.21.576539

**Authors:** José Manuel García-Martínez, Ana Chocarro-Calvo, Javier Martínez-Useros, Nerea Regueira-Acebedo, María Jesús Fernández-Aceñero, Alberto Muñoz, María Jesús Larriba, Custodia García-Jiménez

## Abstract

Cancer initiation and progression result from both genetic alterations and epigenetic reprograming caused by environmental or endogenous factors which can lead to aberrant cell signalling. Most colorectal cancers (CRC) are linked to the abnormal activation of the Wnt/ β-catenin pathway, whose key feature is the accumulation of acetylated β-catenin protein within the nucleus of colon epithelial cells. Nuclear β- catenin acts as a transcriptional co-activator that alters the expression of many target genes involved in cell proliferation and invasion. The most active vitamin D metabolite 1,25-dihydroxyvitamin D3 (1,25(OH)2D3, calcitriol) can antagonize the over-activated Wnt/ β-catenin pathway via binding to its high affinity receptor VDR. Here, we show that the activation of the SIRT1 deacetylase by 1,25(OH)2D3-bound VDR promotes deacetylation and nuclear exclusion of β-catenin and, consequently, the downregulation of its pro-tumorigenic target genes and the inhibition of human colon carcinoma cell proliferation. Notably, orthogonal SIRT1 activation systematically drives nuclear exclusion of β-catenin, highlighting the key role of SIRT1 in CRC. Since nuclear localization of β-catenin is a critical driver of CRC initiation and progression that requires its acetylation, our results provide a mechanistic basis for the epidemiological evidence linking vitamin D deficiency and increased CRC risk and mortality.

## INTRODUCTION

Epidemiological studies suggest that vitamin D (cholecalciferol, VD) deficiency may be a risk factor for developing and dying of cancer, particularly for colon/colorectal cancer (CRC) [1–4]. However, data from supplementation studies in the human population are controversial, and confirmation of a clinically relevant anti-CRC effect of VD in well-designed prospective randomized trials is pending [5–9]. Post-hoc analyses indicate that this discrepancy may rely on the lack of patient stratification in the clinical trials, suggesting that VD intervention is effective in the deficient/insufficient VD subjects but not in VD sufficient individuals, and is also dependent on conditions such as patient body mass, ethnicity, mutational status, or genotype of VD-related genes [6,10,11]. Importantly, many mechanistic experimental studies show a wide range of antitumoral effects of VD in CRC and other neoplasia that strongly support a protective VD action [12–14].

VD is incorporated in humans through diet or is synthesized in the skin by solar ultraviolet radiation-dependent transformation of 7-dehydrocholesterol. The active VD metabolite is 1α,25-dihydroxyvitamin D3 (1,25(OH)2D3, calcitriol, which results from two consecutive hydroxylation of VD, the first in the liver and the second in the kidney or in many epithelial and immune cell types in the organism [1,15]. 1,25(OH)2D3 binds to a member of the nuclear receptor superfamily of transcription factors, the vitamin D receptor (VDR). Upon ligand binding, VDR regulates the pleiotropic actions of VD including its many anticancer effects on cell survival, proliferation and differentiation [1,13,15–17].

The anti-CRC properties of liganded VDR largely rely on its capacity to interfere with a well-recognized CRC driver, the canonical Wnt/β-catenin signalling pathway that regulates many protumoral genes and which is abnormally over-activated in most CRC [18]. Consistently, germline *Vdr* deletion in *Apc*^Min/+^ mice increases Wnt/β- catenin signalling activation and intestinal tumour load [19,20]. From the many possible steps to block Wnt/β-catenin signalling, the most precise is acting directly on β-catenin, its downstream transcriptional effector. Acetylation of β-catenin promotes its nuclear localization and transcriptional activity in cancer cells [21–23]. β-catenin acetylation is increased in CRC cells by the combination of Wnt signalling and enhanced glucose uptake. The coincidence of Wnt and high glucose increases the levels and activity of the acetyl transferase EP300 and inhibits SIRT1-driven deacetylation of β-catenin without changing SIRT1 levels [21] by mechanisms that remain unknown.

Since 1,25(OH)2D3 induces SIRT1 activity in CRC cells [24], we investigated whether 1,25(OH)2D3 impedes the Wnt/β-catenin pathway via regulation of SIRT1. In particular, we examined the intriguing possibility that 1,25(OH)2D3 impacts on CRC by blocking the glucose-driven β-catenin acetylation that enhances Wnt/β-catenin signalling.

## RESULTS

### 1,25(OH)2D3 reverses the acetylation of SIRT1 promoted by Wnt and induces β- catenin nuclear export in CRC cells

1,25(OH)2D3-bound VDR can induce SIRT1 auto-deacetylation and activation [24]. Since SIRT1 targets nuclear exclusion of β-catenin in cancer cells [21] and 1,25(OH)2D3 interferes with Wnt/β-catenin signalling, we first asked whether SIRT1 mediates this effect of 1,25(OH)2D3.

SIRT1 acetylation is a hallmark mark of its inactivity [24]. Given the importance of SIRT1 acetylation for its activity, we studied whether Wnt signals could affect the acetylation status of SIRT1 in nuclear extracts of CRC cells. Exposure of cells to Wnt3A, or LiCl to mimic Wnt signaling, strongly induced SIRT1 acetylation that was reversed by treatment with 1,25(OH)2D3 (Figure 1A). Additionally, 1,25(OH)2D3 increased SIRT1 levels in CRC cells independently of the presence of Wnt signals (Figure 1Β). Thus, 1,25(OH)2D3 ensures SIRT1 induction by increasing both its levels and its deacetylation. Interestingly, co-immunoprecipitation assays demonstrated that 1,25(OH)2D3 favored SIRT1/β-catenin interaction (Figure 1C). Consistently, 1,25(OH)2D3 promoted β-catenin deacetylation in CRC cells (Figure 1D), which paralleled β-catenin nuclear exclusion examined by western blotting (Supplementary Figure S1A) and immunofluorescence analyses (Figure 1E and Supplementary Figure S1Β). Accordingly, 1,25(OH)2D3 reduced the expression of β-catenin target genes associated with tumor progression such as *MYC* and *CCDN1* (Figure 1F, 1G and Supplementary Figure S1C). Notably, 1,25(OH)2D3 could interfere with Wnt signalling independently of whether it was added before or after Wnt activation (Supplementary Figure S1D), suggesting a potential for both prevention and treatment.

**Figure 1.**
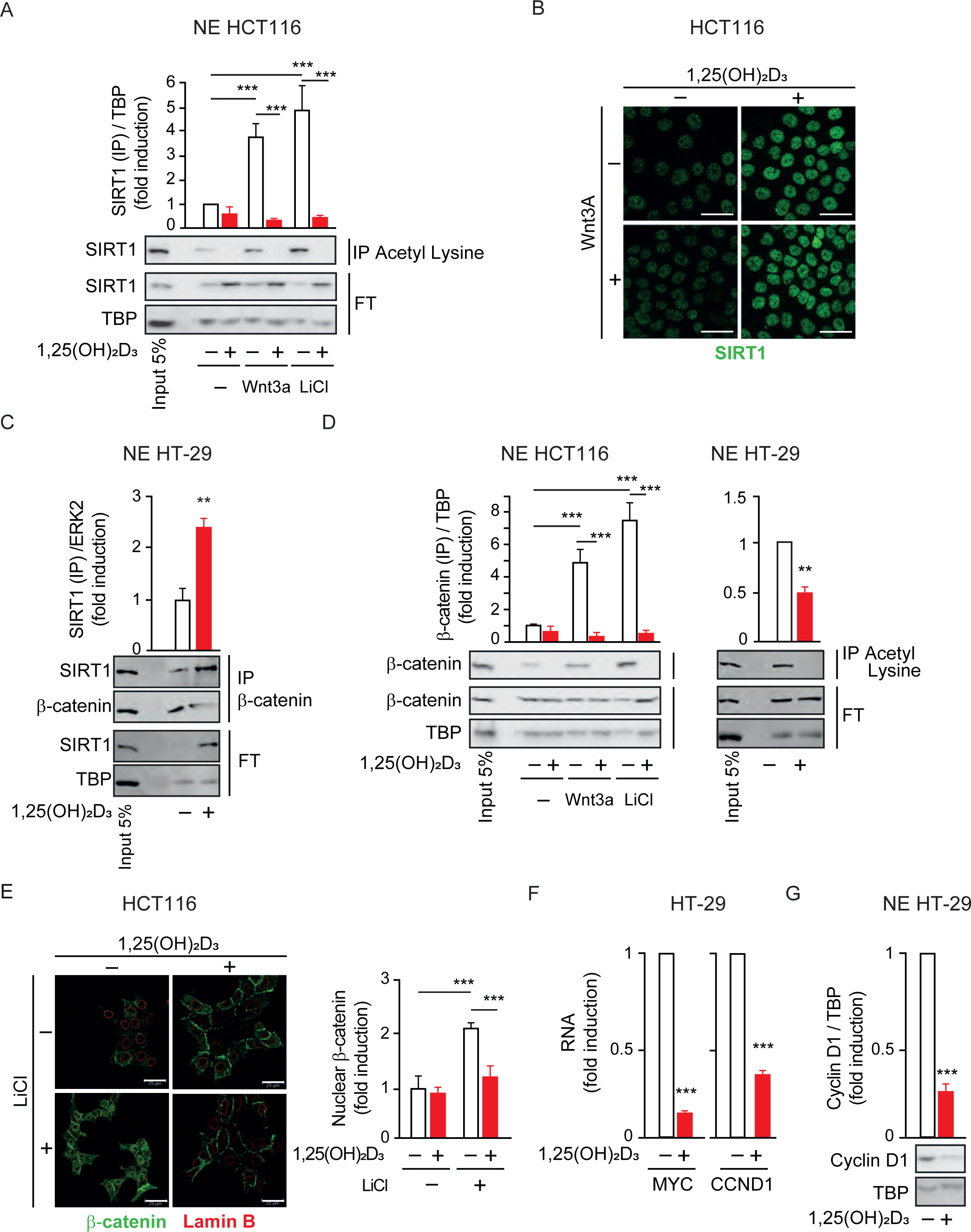
1,25(OH)2D3 Reverses SIRT1 Inactivation by Wnt through β−catenin relocation. (A)-(G) HCT 116 or HT-29 CRC cells cultured under standard conditions and where indicated, treated with Wnt3A (100 ng/ml) or LiCl (40 mM, to mimic Wnt3A) and/or 1α,25-dihydroxyvitamin D3 (1,25(OH)2D3),100 nM, added 24 hours before harvesting. Cell extracts were fractionated and nuclear extracts (NE) of indicated cells were analyzed (A), (C-D) and (G). (A) Western-blot analysis of the effects of Wnt3A, LiCl and/or 1,25(OH)2D3 on the acetylation levels of SIRT1. Nuclear extracts (NE) from HCT 116 CRC cells were immunoprecipitated (IP) using anti-acetyl-lysine antibodies. Representative blots and statistical analysis using TBP in the Flow though (FT) as loading controls. (B) Confocal imaging of the effect of 1,25 (OH)2D3 on SIRT1 levels (green) in HCT 116 CRC cells cultured in the presence/absence of Wnt3A proteins. (C) 1,25(OH)2D3 effects on the SIRT1 / β−catenin interaction in the nuclei of CRC HT-29 cells. Input (5%) and flow through (FT) are shown. ERK2 serves as loading control. (D) Western-blot analysis of the effects of Wnt3A, LiCl and/or 1,25(OH)2D3 on the acetylation levels of nuclear β−catenin. Nuclear extracts (NE) from the indicated CRC cells were immunoprecipitated using anti-acetyl-lysine antibodies. Representative blots and statistical analysis using TBP in the Flow though (FT) as loading controls. (E) Confocal imaging analysis of 1,25 (OH)2D3 effects on nuclear exclusion of β−catenin in HCT 116 cells; nuclear envelope, marked with Lamin B antibody (red) and β−catenin (green). Right panel: Fluorescence intensity was quantified in 3 independent experiments using ImageJ software and for each experiment, 3 different fields were evaluated per slide. Scale bars: 25µm. (F)-(G) analysis of the effect of 1,25 (OH)2D3 on the expression of Wnt-target genes in CRC cells HT-29. (F) RT-qPCR analysis of cyclin D1 (CRCND1) and MYC. Values normalized with endogenous control (18S RNA) are referred as fold induction over cells without 1,25(OH)2D3. (G) western-blot analysis of cyclin D1 protein levels. Representative blots and statistical analysis using TBP as loading control. Statistical analysis by Student t-test of 3 independent experiments; values represent mean ± SEM of triplicates; * p< 0.05; ** p< 0.01; *** p< 0.001.

### SIRT1 activity inversely correlates with the level of nuclear β-catenin during human CRC progression

The striking reversal of the Wnt effect on SIRT1 and β-catenin exerted by 1,25(OH)2D3 in CRC cells led us to investigate whether SIRT1 levels were related to β-catenin localization in human colorectal tumors, in which nuclear accumulation of β-catenin is a marker of poor prognosis. Human tissue microarrays containing samples (n = 109) from primary tumors of stage II and stage IV patients and from liver metastasis of the latter were examined to understand the relationship between SIRT1 level and activity and the subcellular localization of β-catenin. SIRT1 activity was inferred by evaluating the acetylation of its specific well characterized substrate, lysine 9 of histone 3 (AceH3K9).

Although the level of SIRT1 did not change (Figure 2A), AceH3K9 levels, a mark for SIRT1 inactivity increased significantly from primary tumors of stage II patients to metastasis (*P* = 0.027) (Figure 2B). Accordingly, nuclear β-catenin level increased (Figure 2C) while that of cytoplasmic β-catenin was reduced (Figure 2D) through tumor progression as expected.

**Figure 2.**
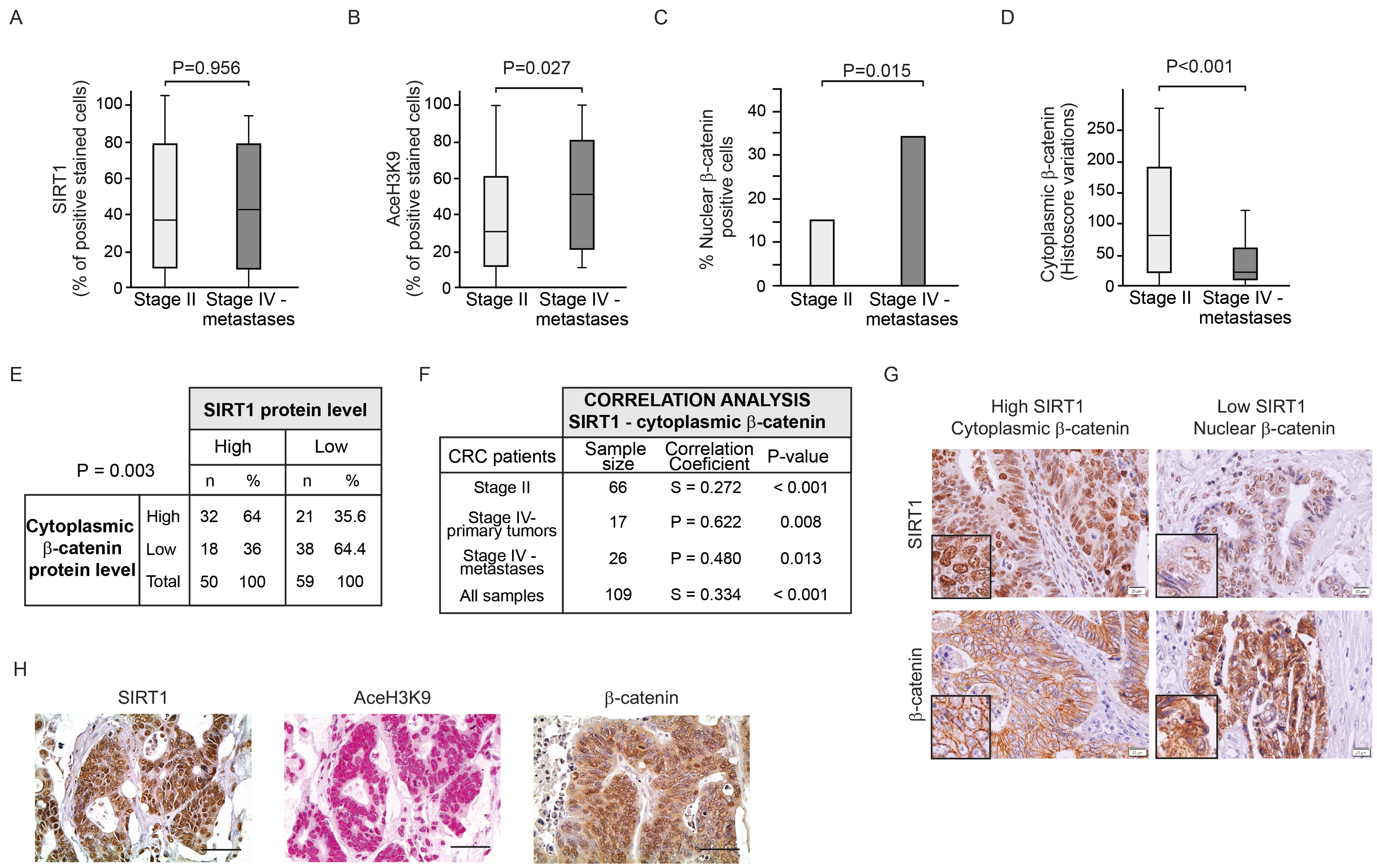
SIRT1 Activity Decline and Nuclear β-catenin Rise in Human CRC Progression Analysis of SIRT1 levels and activity in TMAs of human CRC samples. Box plots for: SIRT1expression (A) and AceH3K9 expression (B), as percentage of positive stained cells, and cytoplasmic β-catenin Hscore expression (C). (D) Bar-graph showing the percentage of CRC human samples with β-catenin nuclear pattern. P<0.05 were considered statistically significant. (E) Association analysis between SIRT1and cytoplasmic protein levels of β-catenin in all CRC samples (primary tumours of stage II & stage IV primary and liver-paired metastases) as described in the methods section under the paragraph *human samples*. Association analysis was performed with Chi-square test. (F) Correlation analysis between cytoplasmic β-catenin and SIRT1 protein levels on CRC patient biopsies of stage II & stage IV primary tumours and their liver-paired metastases. Pearson (P) or Spearman (S) analyses. (G) Representative micrographs at 40X from immunostaining for SIRT1 and its targeted substrates AceH3K9 and β-catenin from the same CRC patient. Scale bars: 20µm. (H) CRC patient samples with high SIRT1 levels show high levels of AceH3K9 and mainly nuclear β-catenin. Representative micrographs at 40X; scale bars: 20µm.

Interestingly, a significant positive association (*P* = 0.003) (Figure 2E) and correlation (R = 0.334, *P* < 0.001) (Figure 2F) between SIRT1 protein levels and cytoplasmic β-catenin levels were found. In addition, significant positive correlations between these parameters were also observed considering separately primary tumors from stage II patients (R = 0.272, *P* < 0.027), primary tumors from stage IV patients (R = 0.622, *P* = 0.008) and liver metastases (R = 0.480, *P* = 0.013) (Figure 2F). Thus, human CRC samples with a low level of SIRT1 tend to have a high level of nuclear β- catenin and *vice versa*. Representative photographs show the coincidence of low SIRT1 level with high nuclear β-catenin in consecutive sections. Samples with high SIRT1 level tend to have high cytoplasmic β-catenin (Figure 2G), although there were samples with high level of SIRT1 and nuclear β-catenin, which could be explained if SIRT1 activity was diminished, as it seems to happen by analysing their high AceH3K9 level (Figure 2H). Collectively, these results suggest that decreased SIRT1 activity (independently of its levels) drives a failure to export β-catenin from the nucleus, which may worsen CRC prognosis.

### SIRT1 is required for nuclear exclusion of β*-*catenin by 1,25(OH)_2_D_3._

To understand whether the nuclear export or accumulation of β-catenin was caused by SIRT1 induction or inhibition, we modulated SIRT1 activity. Treatment with the general sirtuin inhibitor nicotinamide (NAA) prevented the nuclear depletion of β- catenin induced by 1,25(OH)2D3 in CRC cells as analysed by immunofluorescence (Figure 3A) and western blotting (Figure 3Β and Supplementary Figure S2A). Similar results were obtained with the more specific SIRT1 inhibitor EX527 (Figure 3C). SIRT1 was also depleted using several siRNAs (Supplementary Figure S2Β) and its depletion also blocked the 1,25(OH)2D3-mediated nuclear depletion of β-catenin (Figure 3D and Supplementary Figure S2C). In line with these results, SIRT1 inhibition with EX527 abolished the antiproliferative effect of 1,25(OH)2D3 on CRC cells (Figure 3E).

**Figure 3.**
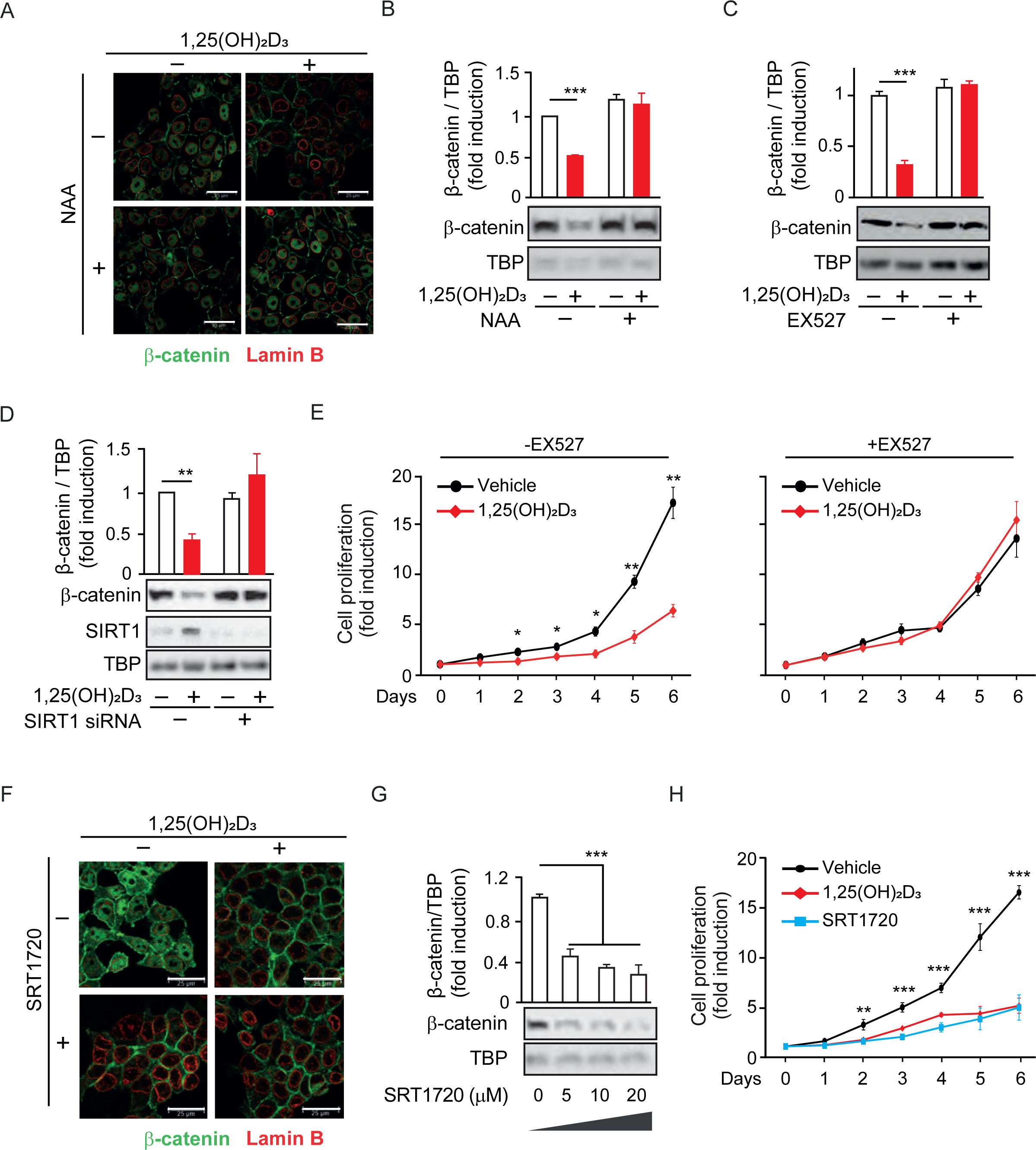
SIRT 1 is required for nuclear exclusion of β−catenin by 1,25(OH)_2_D_3_. HCT 116 CRC cells were cultured with the indicated sirtuin inhibitors or activators and treated or not with 1,25 (OH)2D3 as previously indicated. NAA: Nicotinamide (300 µM), general sirtuin inhibitor. EX527: Selisistat (10 μM), selective SIRT1 inhibitor. SRT1720 (10 μM): selective SIRT1 activator. The effects were analyzed by confocal imaging, scale bars: 25µm. (A), (G); western blotting of nuclear extracts using TBP as loading control (B-D), (H) or proliferation curves (E-F), (I). (A)-(B) Effects of 1,25 (OH)2D3 and NAA on β-catenin subcellular localization. (A) Confocal imaging with nuclear envelope marked with Lamin B antibody (red) and β-catenin (green). (B) Western-blot analysis. Representative blots and statistical analysis. (C) Effect of EX527 on nuclear levels of β-catenin, analysed as in (B). (D) Effect of SIRT1 depletion using a mix of siRNAs from Dharmacon on nuclear levels of β-catenin. (E-F) Effect of EX527 on the 1,25(OH)2D3-driven blockade of HCT 116 CRC cell proliferation. (G) Effect of the SIRT1 activator SRT1720 on subcellular localization of β-catenin analysed by confocal imaging with nuclear envelope marked in red with Lamin B antibody and β-catenin (green). (H) Dose-dependent effect of the specific SIRT1 inducer SRT1720 on nuclear β-catenin levels. (I) Proliferation curves to compare the response to 1,25 (OH)2D3 and SRT1720 in CRC cells. Statistical analysis by Student t-test (B-F) and (I) or ANOVA (H) of 3 independent experiments; values represent mean ± SEM of triplicates; * p< 0.05; ** p< 0.01; *** p< 0.001.

Conversely, the specific activation of SIRT1 by the small molecule SRT1720 drove nuclear exclusion of β-catenin, as shown by immunofluorescence (Figure 3F) or by western blotting (Figure 3G). Similar antiproliferative effects were obtained by activation of SIRT1 with either 1,25(OH)2D3 or SRT1720 (Figure 3H). These results show that the 1,25(OH)2D3-driven depletion of nuclear β-catenin that interferes Wnt signalling is critically mediated by SIRT1 activity.

### A constitutively active SIRT1 mutant mimics the reduction of nuclear β-catenin by 1,25(OH)_2_D_3._

Given the role of SIRT1 on the subcellular localization of β-catenin, we exogenously expressed SIRT1 to ask whether it mediates 1,25(OH)2D3 action controlling β-catenin acetylation and localization. To this end, Myc-tagged SIRT1 wild-type (WT), or the inactive H363Y [25] or the constitutively active K610R [24] SIRT1 mutants were expressed to similar levels in CRC cells. Nuclear extracts from cells transfected with SIRT1 WT or mutants were immunoprecipitated using anti-acetyl-lysine antibodies and subjected to western blotting to evaluate the extent of β-catenin acetylation. In the nuclei of CRC cells, acetylated β-catenin decreased upon expression of WT SIRT1, increased after expression of the inactive H363Y mutant and was lost upon expression of the constitutively active K610R mutant (Figure 4A). Consistently, the nuclear levels of β-catenin reflected its acetylation status and was reduced by half after expression of WT SIRT1, whereas the inactive H363Y mutant had no effect, and nuclear β-catenin was almost completely depleted (4-5-fold reduction) following expression of the constitutively active K610R SIRT1 (Figure 4Β).

**Figure 4.**
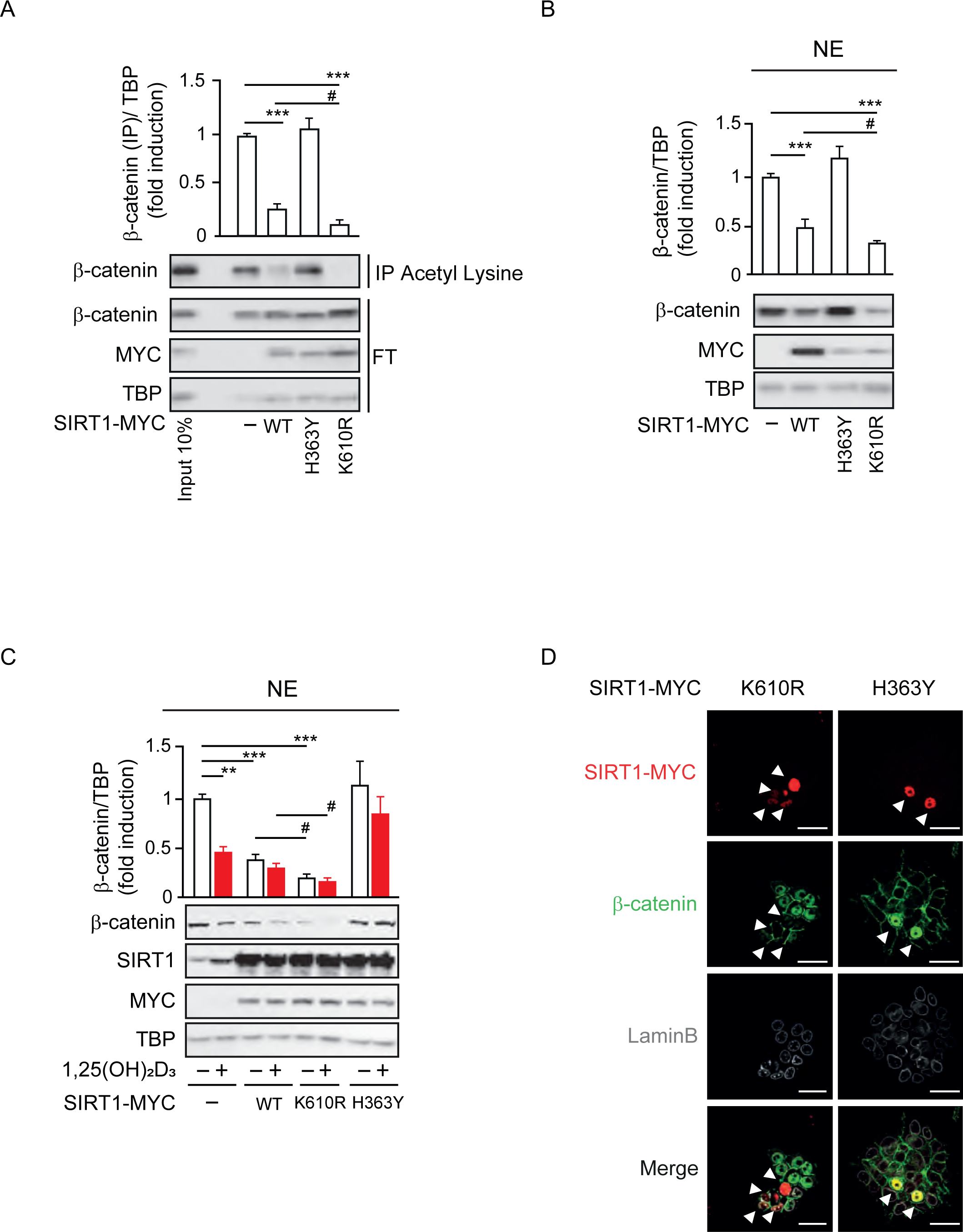
Constitutively active SIRT1 mutant exclude β−catenin from the nucleus as 1,25(OH)_2_D_3_. HCT 116 CRC cells cultured in DMEM were transiently transfected with pcDNA expression vectors: empty (-), myc tagged SIRT1 wild type (WT) or mutants: H363Y inactive or K610R, 48 h before harvesting. LiCl was used at 40 mM and 1,25 (OH)2D3 (100 nM) was added where indicated for the last 24 h. Extracts were fractionated. (A-C) Western blot analysis of nuclear extracts (NE) using TBP as loading control and MYC as expression control; representative western-blots and statistical analysis. (D) Confocal imaging, scale bars represent 25 μm. (A) Effect of SIRT1 expression, wild type (WT) or indicated activity SIRT1 mutants (inactive H363Y or active K610R) on the acetylation levels of nuclear β-catenin analysed after immunoprecipitation (IP) with anti-acetyl lysine antibodies; FT: Flow through. (B) Effect of SIRT1 expression (WT and mutants) on nuclear levels of β-catenin. (C) Interference by 1,25 (OH)2D3 of the effect of SIRT1 expression (WT and mutants) on nuclear β-catenin levels. (D) Confocal imaging of the myc-tagged SIRT1 mutant (red), β-catenin (green), Lamin B to mark the nuclear envelope (white). Merge panels at the bottom. Statistical analysis by One Way ANOVA of n=3 independent experiments. Values represent mean ± SEM; *p<0.05; **p<0.01; *** p<0.001.

We also examined β-catenin subcellular localization in response to 1,25(OH)2D3 upon expression of these mutants. 1,25(OH)2D3 treatment in control cells reduced the level of nuclear β-catenin up to 50%, a small reduction was measured in cells expressing WT SIRT1, whereas in cells expressing the constitutively active K610R SIRT1 nuclear β-catenin maximally diminished by 75% regardless of 1,25(OH)2D3 treatment (Figure 4C). Conversely, cells expressing the inactive H363Y SIRT1 mutant exhibited higher basal nuclear levels of β-catenin and were also unresponsive to 1,25(OH)2D3 (Figure 4C). Of note, immunofluorescence analyses confirmed that K610R SIRT1 was constitutively active and depleted β-catenin from the nucleus of HCT 116 cells while, in contrast, the inactive H363Y SIRT1 mutant led to nuclear accumulation of β-catenin (Figure 4D). Collectively, our data indicated that 1,25(OH)2D3 induces SIRT1 activity, which controls β-catenin acetylation, subcellular localization, and transcriptional activity.

### SIRT1 activity offers a surrogate target to inhibit the Wnt/β-catenin pathway in *cases of 1,25(OH)_2_D_3_ unresponsiveness*

VDR expression is the main determinant of cell responsiveness to 1,25(OH)2D3 and is often downregulated in advanced CRC, which together with frequent VD deficiency implies that these patients would probably not benefit from the anticancer effects of 1,25(OH)2D3 [20,26–28]. We sought to investigate the potential of targeting SIRT1 under these conditions by using HCT 116 cells in which VDR expression was downregulated by means of shRNA[19]. Western blotting and immunofluorescence analyses showed that the nuclear content of β-catenin was slightly higher in ShVDR cells, which did not respond to 1,25(OH)2D3 by excluding β-catenin from their nuclei as expected. In contrast, ShControl cells responded to 1,25(OH)2D3 with strong nuclear β-catenin depletion (Figure 5A-B).

**Figure 5.**
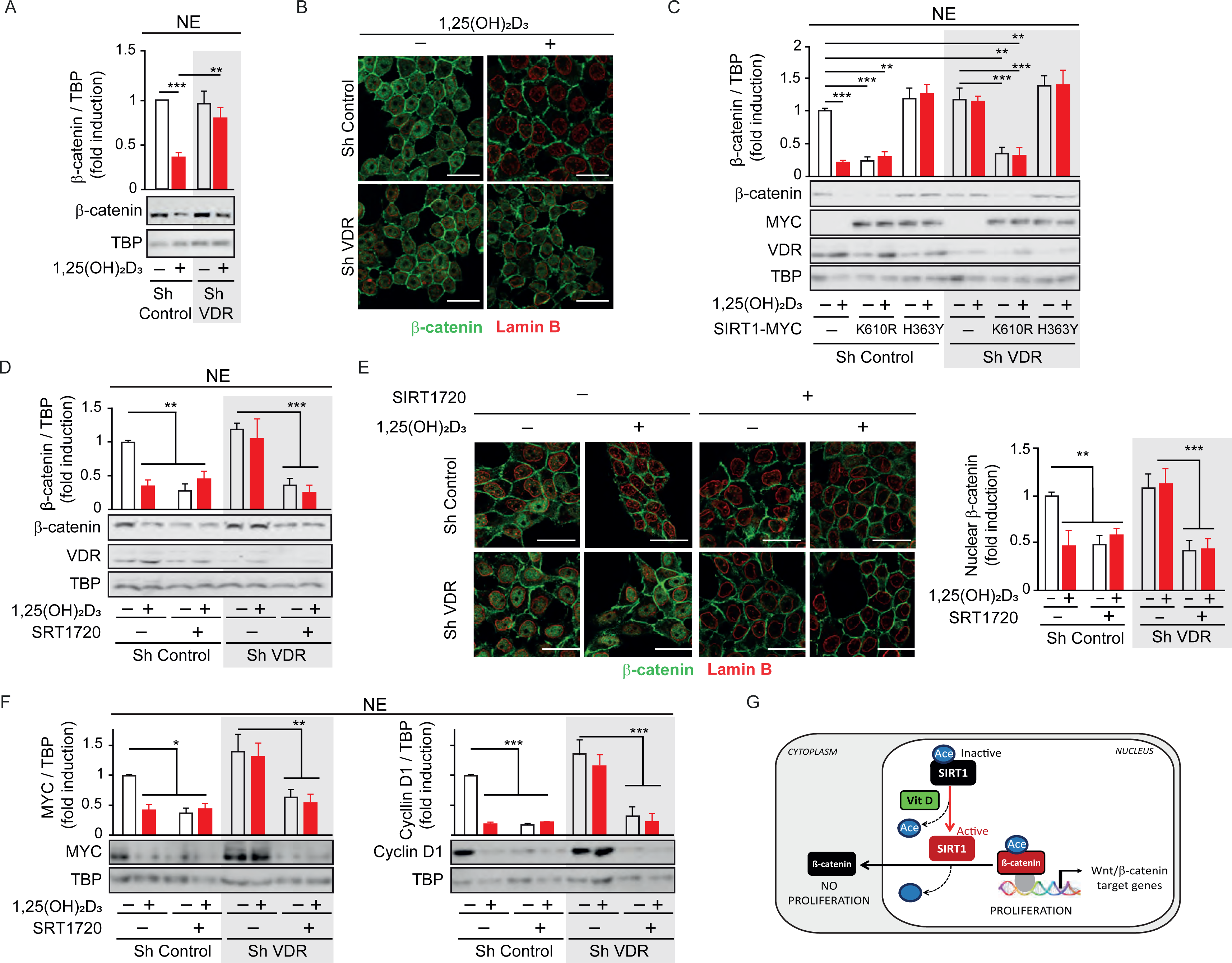
SIRT1 activity offers a surrogate target in vitamin D unresponsive and SIRT1 expressing CRC. HCT 116 derived cells were obtained by stable knock-down of Vitamin D receptor (VDR) using specific shRNA targeting VDR to obtain Sh VDR cells, that were compared to ShControl cells that contain normal VDR levels [19]. Cells were cultured in DMEM under standard conditions, treated with 1,25 (OH)2D3 and fractionated as previously described; nuclear extracts were analyzed. For Panel C cells were transfected to express SIRT1 mutants H363Y and K610R to comparable levels or the empty vector (Ø). (A) Western blot analysis of nuclear β-catenin response to 1,25 (OH)2D3 in cells previously stimulated with LiCl and depleted or not of VDR. (B) Effect of 1,25 (OH)2D3 on β-catenin subcellular localization in cells previously stimulated with LiCl. Lamin B (red) marks the nuclear envelop to show nuclear accumulation or exclusion of β-catenin (green) in response to 1,25 (OH)2D3. Scale bars represent 25 μm. (C) Nuclear β-catenin response to 1,25 (OH)2D3 in Sh VDR cells compared to Sh Control cells, upon expression of the SIRT1 activity mutants H363Y or K610R. Levels of expression of mutants are detected with anti-MYC antibody. VDR levels are shown for depletion control. TBP as loading control. (D) Nuclear β-catenin responses to 1,25 (OH)2D3 or to the SIRT1 activator SRT1720 in Sh VDR and Sh Control cells. Representative western blots and statistical analysis. TBP as loading control and VDR levels as depletion control. (E) Confocal imaging of the nuclear β-catenin (green) response to 1,25 (OH)2D3 or the SIRT1 activator SRT1720 in Sh Control and Sh VDR cells. Lamin B at the nuclear envelop is shown in red. Scale bars represent 25 μm. (F) Nuclear levels of MYC AND Cyclin D1 in response to 1,25 (OH)2D3 or to the SIRT1 activator SRT1720 in Sh VDR and Sh Control cells. Representative western blots and statistical analysis. TBP as loading control. (G) Scheme representative of the new 1,25 (OH)2D3 driven mechanism to block cancer cell proliferation through nuclear exclusion of β-catenin Statistical analysis by One-Way ANOVA of 3 independent experiments; values represent mean ± SEM of triplicates; * p< 0.05; ** p< 0.01; *** p< 0.001.

Next, we examined the potential to overcome the lack of response to 1,25(OH)2D3 in ShVDR HCT116 cells by expressing the SIRT1 mutants (Figure 5C). Interestingly, expression of the constitutively active K610R SIRT1 mutant reduced nuclear β-catenin levels in both ShControl and ShVDR cells to an extent similar to that achieved by 1,25(OH)2D3 treatment in ShControl cells. As expected, the inactive H363Y SIRT1 did not modify nuclear β-catenin levels (Figure 5C). These results suggested that targeting SIRT1 activity might alleviate the effects of VD deficiency or unresponsiveness. Indeed, the specific SIRT1 activator SRT1720 reduced nuclear β-catenin levels in both ShControl and ShVDR cells, as shown in nuclear extracts by western blotting (Figure 5D) or in whole cells by immunofluorescence analysis (Figure 5E). Accordingly, SRT1720 reduced the levels of Wnt target proteins such as MYC and Cyclin D1 in ShControl cells at a comparable magnitude to 1,25(OH)_2_D_3._ Notably, SRT1720 also reduced MYC and Cyclin D1 protein levels in the vitamin D unresponsive ShVDR cells (Figure 5F).

These results indicate that Wnt/β-catenin signalling can be reduced by targeting SIRT1 in cells unresponsive to 1,25(OH)2D3. Altogether, this work demonstrates that 1,25(OH)2D3 (i) induces reversal of SIRT1 acetylation imposed by Wnt, (ii) favors SIRT1/β-catenin interaction, (iii) increases β-catenin deacetylation, (iv) promotes nuclear exclusion of β-catenin and (v) decreases expression of key Wnt-target genes (Figure 5G).

## DISCUSSION

Here we reveal that SIRT1 activity is a critical mediator of the anticancer properties of 1,25(OH)2D3 related to its interference of Wnt/β-catenin signalling in CRC. The critical importance of Wnt signalling in CRC is emphasized by the fact that over 94% of primary and up to 96% of metastatic colorectal tumors contain mutations that aberrantly activate Wnt/β-catenin signaling [29,30] Importantly, Wnt/β-catenin induces a gene signature (largely coincident with that of intestinal stem cells) that identifies CRC stem cells and predicts disease relapse [31]

Previous work of our group and others has unveiled 1,25(OH)2D3 targets that interfere Wnt/β-catenin signalling, such as VDR-dependent increased E-cadherin expression and improved E-cadherin-β-catenin interactions at the membrane [32], competition between VDR and Tcf/Lef transcription factors to bind β-catenin [12,13,33], and the induction of the Wnt inhibitor DKK-1 [34]. This work demonstrates that inhibition or depletion of SIRT1 leaves 1,25(OH)2D3 unable to interfere Wnt target gene expression or proliferation in CRC cells, thus providing a mechanism that mediates previously described effects.

Surely, acetylation governs protein-protein and protein-DNA interactions due to increased size of the lysine side chain and neutralization of its positive charge, potentially disrupting salt bridges and introducing steric bulks [35–37]. Indeed, VDR deacetylation by SIRT1 increases its transcriptional activity on specific genes [38] and may as well drive increased expression of alternative VDR target genes capable to interfere Wnt signalling such as E-cadherin or DKK1. In fact, we observe E-cadherin protein levels regulated through SIRT1 modulation. Moreover, SIRT1-driven deacetylation of β-catenin may establish the mark for its preference to bind VDR instead of Tcf/Lef in the nucleus, its capacity to be exported and its interactions with E-cadherin at the plasma membrane. Not surprisingly, there is an increasing interest on the effects of vitamin D on SIRT1 [39], and on the modulation by 1,25(OH)2D3 of VDR and SIRT1 interactions with Fox O and NF-κB as mediators of vitamin D pleiotropism [40,41].

The role of SIRT1 in cancer is controversial and complicated given the multiplicity of its interactions with histones, transcriptional regulators and enzymes and its control of genome stability, cellular differentiation, growth, and metabolism [42,43]. Although[44] reported that SIRT1 inhibition reduces tumorigenesis in animals, suggesting that SIRT1 acts as tumor promoter, [45] reported that SIRT1-deficient embryos exhibit genomic instability and [46] showed that SIRT1 suppressed tumor growth in a mouse colon cancer model driven by Wnt/β-catenin. This last model suggests that the tumor suppressive action of SIRT1 must be exerted on Wnt/β-catenin. Notably, SIRT1 levels and activity are not regulated in parallel and there are CRCs with high SIRT1 levels and low SIRT1 activity [24]. Therefore, reports on increased SIRT1 levels through tumor progression, for example [47], might demonstrate that SIRT1 activity also increases. This is also important to interpret the correlation between SIRT1 overexpression in xenografts and increased tumor size since increased SIRT1 levels may not correlate with increased SIRT1 activity.

Importantly, by using the specific SIRT1 activator SRT1720, the SIRT1 inhibitors NAA and EX527, several siRNAs against SIRT1, and overexpression of SIRT1 WT and constitutive active K610R and inactive H363Y mutants, we convincingly show that the depletion of nuclear β-catenin and the antiproliferative effects exerted by 1,25(OH)2D3 in CRC cells rely on SIRT1 induction and subsequent β-catenin deacetylation. Notably, we also show that the reported inactivation of SIRT1 by Wnt signalling [21,22] correlates with SIRT1 acetylation that is reversed by 1,25(OH)2D3 treatment, reported to activate SIRT1[24].

Increased nuclear β-catenin in HeLa cells upon SIRT1 overexpression [48], led to conclude that SIRT1 deacetylation of β-catenin promotes its nuclear entry. This is at odds with the opposite phenotype reported in this paper and with the established general assumption that β-catenin acetylation improves its stability by inhibiting its ubiquitin-mediated degradation and promotes its nuclear translocation [21,49–52]. Nonetheless, HeLa cells under basal conditions showed abundant SIRT1 with most β- catenin in the cytosol and although nude mice with HeLa xenografts overexpressing SIRT1 developed bigger tumors, this could be achieved independently of Wnt signalling given the pleiotropic effects of SIRT1 on genome stability, metabolism, and general transcription. In addition, SIRT1 acts at many levels on Wnt signalling including reduced expression of Dvl [53] and silencing Wnt antagonists such as DACT1 or sFRPs [54]. In contrast to these studies, our work shows unequivocally that cells expressing constitutively active SIRT1 K610R mutant exhibit nuclei without β- catenin, whereas cells expressing the inactive H363Y mutant accumulate nuclear β- catenin. In addition, we show nuclear β-catenin depletion and deacetylation under SIRT1 activation or over expression.

Importantly, several acetylation sites present in β-catenin (including K19, K49, K345, K354) and targeted by different acetyl transferases (CΒP, EP300, pCAF), govern its affinity for different proteins and may do so in a cell type specific manner. This may offer an explanation to the distinct outcomes of β-catenin deacetylation in distinct cell types. Indeed, acetylation of β-catenin K354 was previously reported to regulate cytoplasm-nuclear shuttling of β-catenin in CRC cells [21]; however, studies that suggest SIRT1 driven nuclear localization of β-catenin in other cells by deacetylation are based on exogenous expression of β-catenin K49 mutants.

Our results show that activation of SIRT1 by 1,25(OH)2D3 reverses the effect of Wnt, excluding β-catenin from the nuclei of CRC cells. VD deficiency is strongly associated epidemiologically to CRC but also to diabetes [12,13,55]. Since the effects described here reverse the potentiation of Wnt signalling exerted by high glucose in diabetes [22,56–60], our results suggest that VD deficiency may be one underlying cause connecting these two diseases.

Wnt signalling regulates development and drives adult tissue stem cell maintenance. Although its malfunction is involved in cancer and many other diseases, it is therapeutically underexploited due to toxicity and limited efficacy [61,62]. Unfortunately, despite the potential of 1,25(OH)2D3 to interfere Wnt/β-catenin signalling, advanced CRCs often loss VDR expression and become unresponsive. Here, we highlight that, downstream of VDR, small molecule activators for SIRT1 such as SRT1720 offer an alternative approach and a therapeutic hope for these cancers if they could be directed specifically to cancer cells. Our results point to SIRT1 as a critical effector of 1,25(OH)2D3 and rise the hope that acting on SIRT1 may circumvent 1,25(OH)2D3 unresponsiveness derived from VD deficiency or VDR loss as it is common in advanced CRC.

## MATERIALS AND METHODS

**Table 1.**
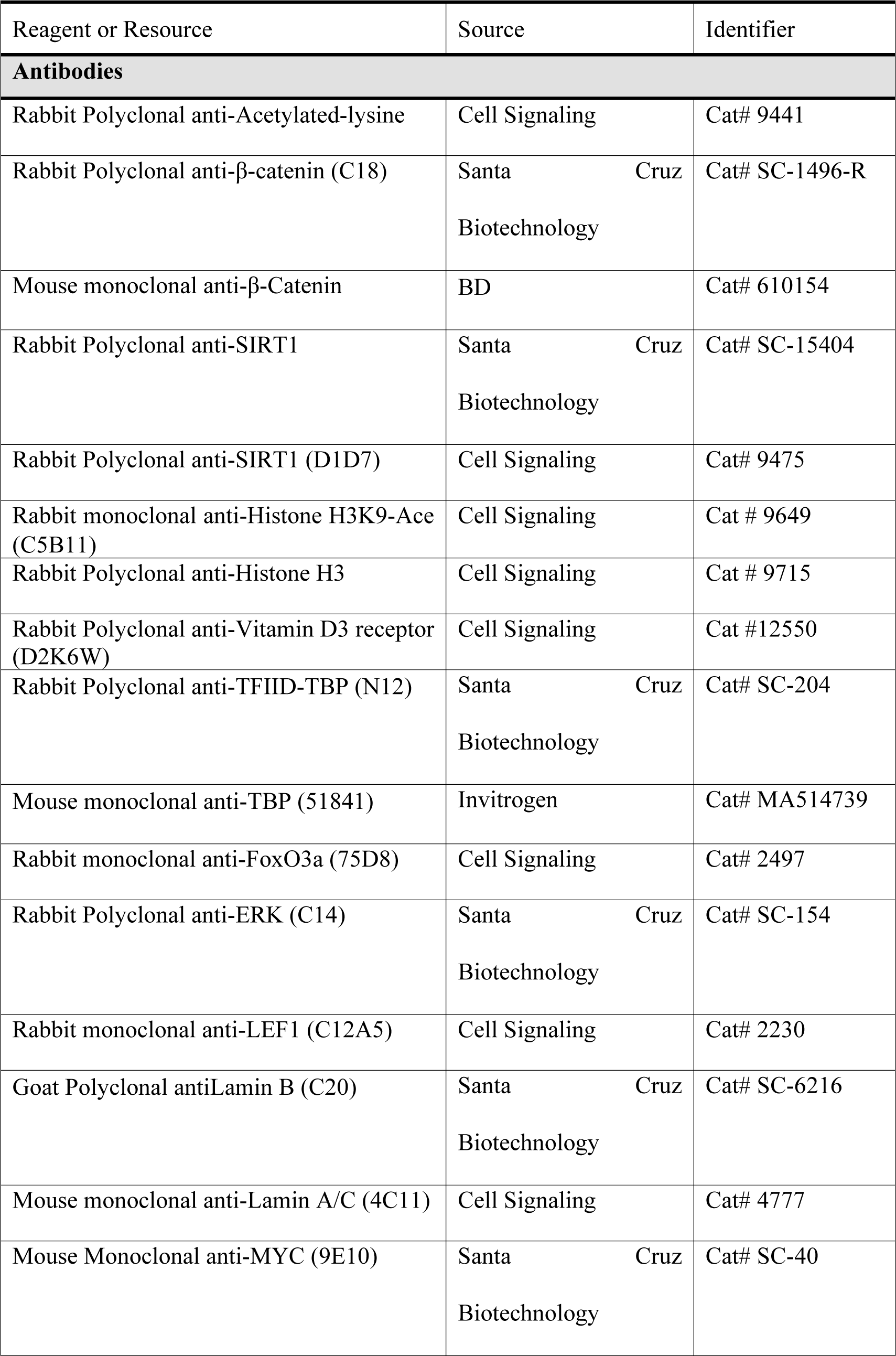
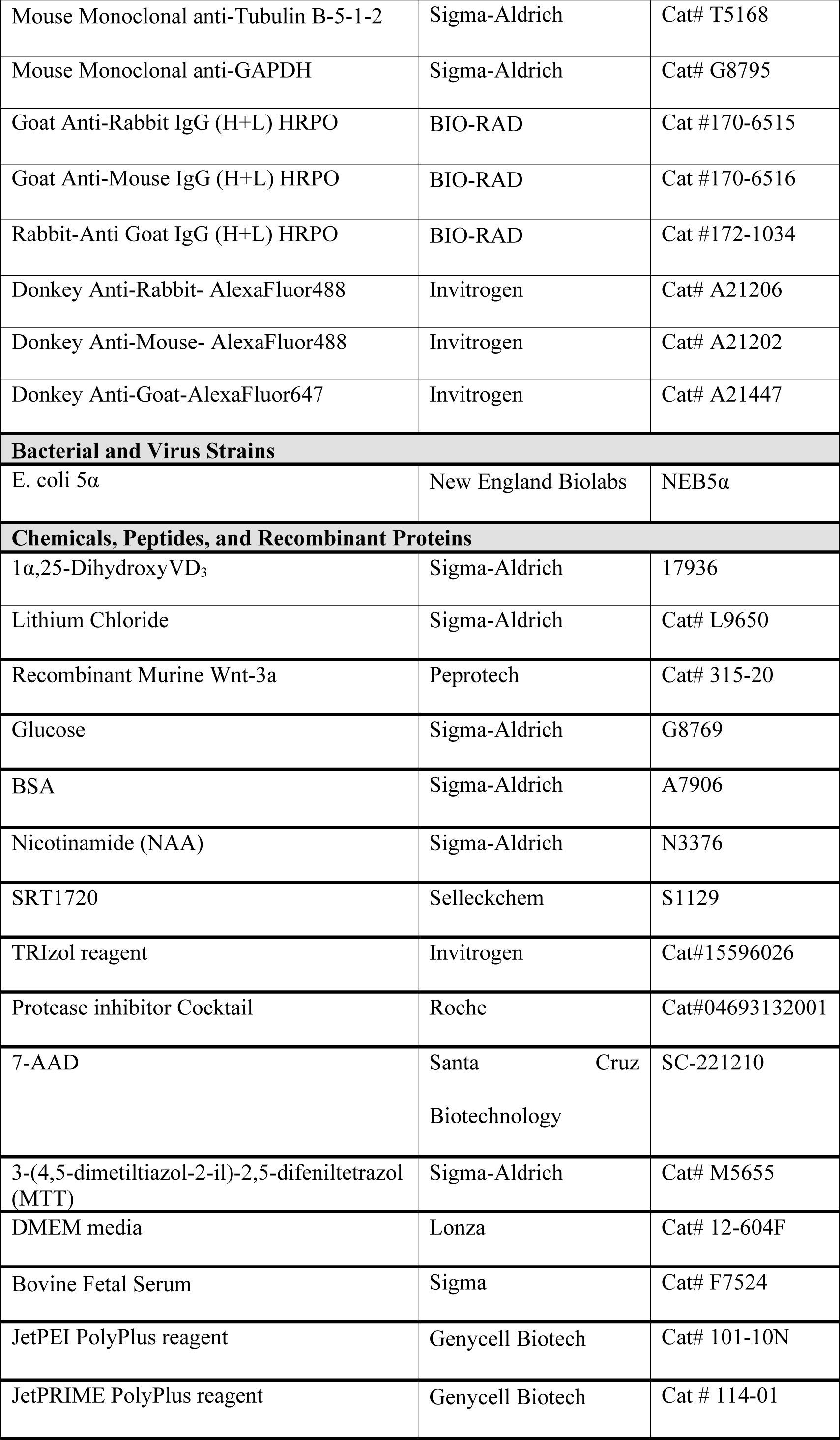
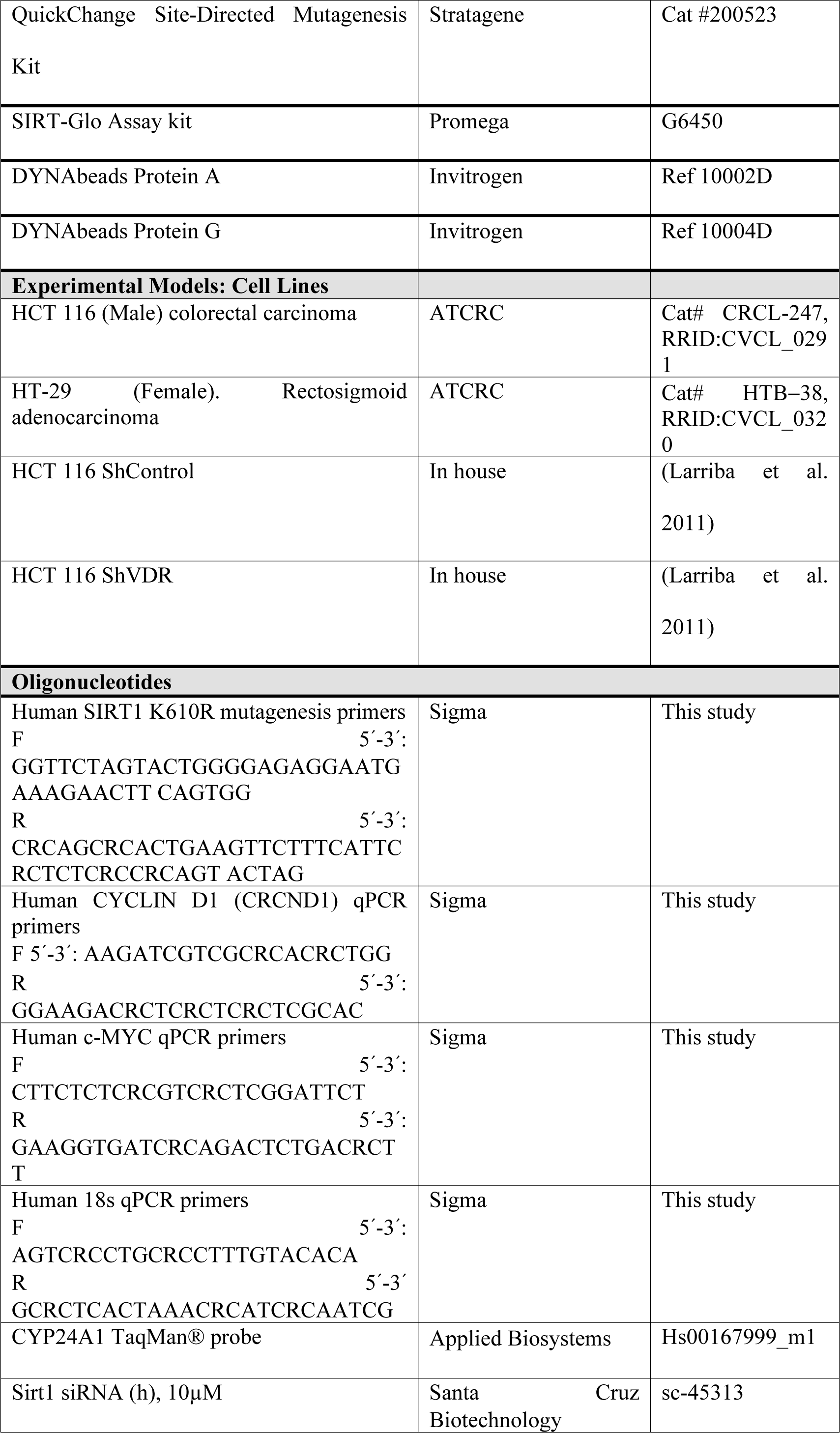
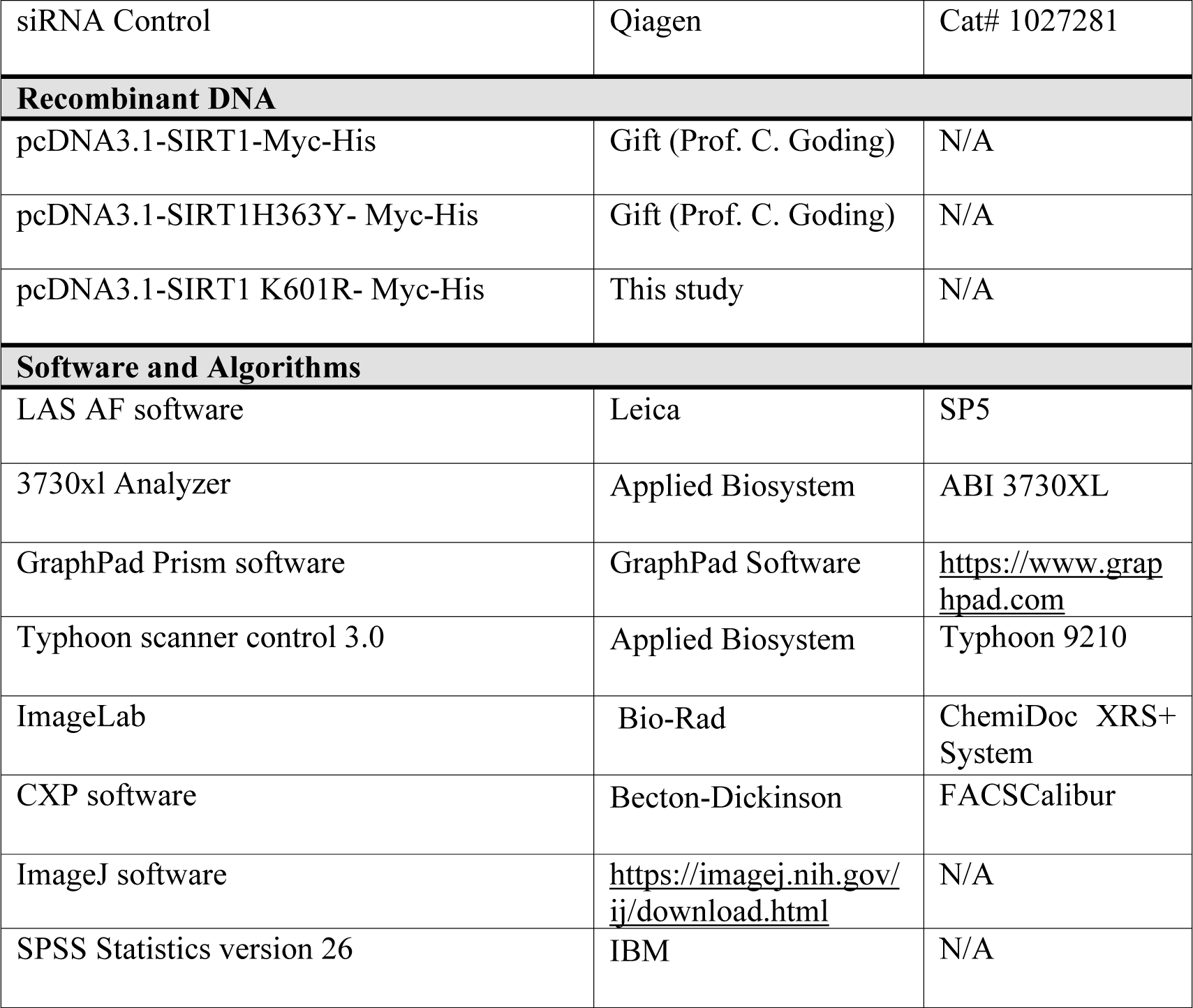
Key Materials & Resources-Key resources.

## METHOD DETAILS

### Colorectal cell panels

Human colorectal adenocarcinoma HT-29 and HCT 116 and HCT 116 derived ShControl and ShVDR cells were cultured in 5%CO2 at 37°C with DMEM containing 25 mM glucose (unless specifically indicated) supplemented with 10% foetal bovine serum (FBS) and 1% penicillin-streptomycin. Cells were treated as indicated for 24h. HCT 116 Sh Control and ShVDR cell lines were derived previously [19]

### Transient transfections

For plasmid transfection, cells were seeded in plates at 50% confluence using JetPei PolyPlus reagent, following the manufacturer’s instructions. After 24h cells were treated as indicated for 24 hours.

For Sirt1 siRNAs, cells plated in six well plates at 50% confluence were transfected with JetPRIME following the manufacturer’s instructions. After 2 days cells were treated 24h with LiCl 40 mM and then another 24h with 1,25(OH)_2_D_3_ before being collected to analyse by western blot.

### Preparation of cell extracts

#### Whole cell extracts

Cells were washed with iced PBS before extract preparation and scraped in RIPA buffer (10 mM Tris HCl [pH 7.4], 5 mM EDTA, 5 mM EGTA, 1% Tryton X100, 10 mM Na_4_P_2_O_7_ [pH 7.4], 10 mM NaF, 130 mM NaCl, 0.1% SDS, 0,5% Na-deoxycholate). After 5 min on ice, cells were pelleted (12000 rpm for 5 min, 4°C) and the supernatant was directly used as whole cell extract or frozen at ™80 °C.

#### Fractionated cell extracts

After washing as before, cells were scraped in hypotonic buffer (20 mM Hepes, [pH 8.0], 10 mM KCl, 0,15 mM EDTA, 0,15 mM EGTA, 0,05% NP40 and protease inhibitors) and incubated on ice for 10 min before adding 1:2 vol of sucrose buffer (50 mM Hepes [pH=8.0], 0.25 mM EDTA, 10 mM KCl, 70% sucrose). Lysates were fractionated (5000 rpm for 5 min at 4°C) to obtain the cytoplasmic fraction in the supernatant. Nuclear pellets were washed twice with washing buffer (20 mM Hepes [pH 8.0], 50 mM NaCl, MgCl_2_ 1.5 mM, 0,25 mM EDTA, 0,15 mM EGTA, 25% glycerol and protease inhibitors), pelleted at 5000 rpm, 5 min at 4°C and resuspended in nuclear extraction buffer (20 mM Hepes[pH 8.0], 450 mM NaCl, MgCl_2_ 1.5 mM, 0,25 mM EDTA, 0,15 mM EGTA, 0,05% NP40, 25% glycerol and protease inhibitors) before centrifugation at 12,000 rpm for 5 min at 4°C to pellet and discard cell debris. The supernatants were used as nuclear fractions.

### Immunoprecipitation

For immunoprecipitation from fractionated extracts the hypotonic buffer was modified by adding 100mM NaCl and 0.1% NP40. For immunocomplex formation, protein A/G-coated magnetic beads (Invitrogen) were washed 3 times with the extraction buffer before coating with the primary antibody for 2 hr at 4 °C in a rotating wheel, followed by 2 washes with the same buffer to eliminates unbound antibody and then extracts are added O/N at 4°C in the rotating wheel. Immunocomplexes were washed twice and used for western blotting.

### Western blotting

Proteins from lysed cells or immunoprecipitates were denatured and loaded on sodium dodecyl sulfate polyacrylamide gels and then transferred to polyvinylidene difluoride membranes (Bio-Rad). After blocking with 5% (w/v) BSA or milk the membrane was incubated with the corresponding primary and secondary antibodies (Bio-Rad). The specific bands were analyzed using Thyphoon or ChemiDoc Imaging Systems (Bio-Rad).

### Immunoflorescence

Cells in cover slips were washed three times and fixed with 4% paraformaldehyde in PBS [pH 7.4] for 10min; washed again; permeabilized (PBS [pH7.4], 0.5% Triton X-100, 0.2% BSA) for 5min; blocked (PBS [pH7,4, 0,05% Triton X-100, 5% BSA) for 1h at room temperature; incubated with primary antibody over night at 4°C, washed three times for 5 min and incubated with the secondary antibody for 1h at room temperature. Slides were mounted and images were acquired using a SP5 confocal microscope (Leica) with a 63x objective. Fluorescence intensity was quantified using Image J software. For each experiment, 3 different fields were evaluated per slide.

### Cell-growth curves

Cell growth was determined by colorimetry. For that, cells were seeded at a density of 20,000 cells per well in a Corning 12-well plate and treated as indicated for 1 to 6 days. After that cells were treated with 3-(4,5-dimethylthiazol-2-yl)-2,5-diphenyltetrazolium bromide (MTT) (Sigma-Aldrich) in a 1:10 ratio in a culture medium and incubated at 37 °C for 3 h. In living cells, MTT is reduced to formazan, which has a purple color. The medium was removed, and the formazan was resuspended in dimethylsulfoxide (DMSO), transferred to p96 plates and analyzed by a Spectra FLUOR (Tecan) at 542 nm. Viability was measured in four independent experiments and then duplicated.

### RT-qPCR

Total RNA was extracted from 3 replicates of colorectal cells using TRIzol reagent (Invitrogen). Reverse transcription of 1μg of RNA was performed according to the manufacturer’s instructions Reagents and detection systems were from Applied biosystems. 18S ribosomal RNA primers served as a nonregulated control. Relative expression was calculated using the Ct method, expressed as 2^−ΔΔCt^ [63]. The PCR efficiency was approximately 100%.

### Human samples

Patient samples were derived from surgical removal at (i) Fundación Jimenez Diaz University Hospital, General and Digestive Tract Surgery Department: 95 patient samples diagnosed with stage II CRC; (ii) at Hospital Clínico San Carlos (HUCSC) 56 primary samples diagnosed with stage IV CRC, and 54 paired liver metastases. The Institutional Review Board (IRB) of the Fundación Jimenez Diaz Hospital, reviewed and approved the study, granting approval on December 9, 2014, by the act number 17/14. All patients gave written informed consent for the use of their biological samples for research purposes. Fundamental ethical principles and rights promoted by Spain (LOPD 15/1999) and the European Union EU (2000/C364/01) were followed. In addition, all patients’ data were processed according to the Declaration of Helsinki (last revision 2013) and Spanish National Biomedical Research Law (14/2007, of July 3).

### TMA, immunohistochemistry, and quantification

Tissue microarrays (TMA) with stage II and stage IV (primary tumors and their paired liver metastases) containing 95, 56 and 54 cores respectively, were constructed using the MTA-1 tissue arrayer (Beecher Instruments, Sun Prairie) for immunohistochemistry analysis. Each core (diameter 0.6 mm) was punched from pre-selected tumor regions in paraffin-embedded tissues. We chose central areas from the tumor, avoiding foci of necrosis. Staining was conducted in 2-μm sections. Slides were deparaffinized by incubation at 60°C for 10 min and incubated with PT-Link (Dako, Agilent) for 20 min at 95°C in low pH to detect SIRT1, or high pH to detect β-Catenin antigens. To block endogenous peroxidase, holders were incubated with peroxidase blocking reagent (Dako, Agilent) and then with the following dilutions of antibodies: anti-SIRT1 (1:50) overnight at 4°C, and anti-β-Catenin (1:500) for 20 min at room temperature. All previously described antibodies presented high specificity. After that, slides were incubated for 20 min with the appropriate anti-Ig horseradish peroxidase-conjugated polymer (EnVision, Dako, Agilent). Sections were then visualized with 3,3’-diaminobenzidine (Dako, Agilent) as a chromogen for SIRT1 and β-catenin or with the chromogen HRP Magenta (Dako, Agilent) for AceH3K9, for 5 min and counterstained with Harrys’ Haematoxylin (Sigma Aldrich, Merck). Photographs were taken with a stereo microscope (Leica DMi1). According to the human protein atlas (available at http://www.proteinatlas.org), a human testis tissue was used as a positive control for anti-SIRT1. Immunoreactivity was quantified blind with an Histoscore (H score) that considers both the intensity and percentage of cells stained for each intensity (low, medium, or high) following this algorithm (range 0–300): H score = (low%) × 1 + (medium%) × 2 + (high %) × 3. Quantification for each patient biopsy was calculated blindly by 2 investigators (MJFA and JMU). AceH3K9 and SIRT1 showed nuclear staining, whereas β-catenin was in the nucleus and cytoplasm. Clinicopathological characteristics of patients are summarized in Table 2.

**Table 2.** Clinicopathological characteristics of CRC patients included in the study.

| Patients Characteristic | N (%) |  |
| --- | --- | --- |
|  | Stage II | Stage IV |
| <u>Gender</u> |  |  |
| Male | 57 (60%) | 27 (48%) |
| Female | 38 (40%) | 23 (41%) |
| N/A | 0 | 6 (11%) |
| <u>Tumor Location</u> |  |  |
| Cecum | 13 (14%) | 3 (5%) |
| Right | 25 (26%) | 10 (18%) |
| Transverse | 7 (8%) | 3 (5%) |
| Left | 5 (5%) | 2 (4%) |
| Sigma | 24 (25%) | 21 (38%) |
| Rectum | 21 (21%) | 12 (21%) |
| N/A | 0 | 5 (9%) |
| <u>Grade Primary Tumor</u> |  |  |
| Well differentiated | 18 (19%) | 15 (27%) |
| Moderately differentiated | 69 (73%) | 27 (48%) |
| Poorly differentiated | 8 (8%) | 5 (9%) |
| N/A | 0 | 9 (16%) |
| <u>Paired Liver Metastasis</u> |  | 54 (96%) |
| N/A |  | 2 (4%) |
| <u>Grade Metastasis Tumor</u> |  |  |
| Well differentiated |  | 12 (21%) |
| Moderately differentiated |  | 29 (52%) |
| Poorly differentiated |  | 4 (7%) |
| N/A |  | 11 (20%) |
| <u>Metastasis at diagnosis</u> |  |  |
| Synchronous |  | 35 (62%) |
| Metachronous |  | 21 (38%) |
| <u>Pt</u> |  |  |
| T1 | 3 (3%) | 2 (4%) |
| T2 | 30 (32%) | 5 (9%) |
| T3 | 61 (64%) | 42 (75%) |
| T4 | 1 (1%) | 4 (7%) |
| N/A |  | 3 (5%) |
| <u>pN</u> |  |  |
| N0 | 95 (100%) | 27 (48%) |
| N1 |  | 16 (29%) |
| N2 |  | 9 (16%) |
| N3 |  | 1 (2%) |
| N/A |  | 3 (5%) |

### Statistical analyses

To determine whether each of the antigens evaluated here were well-modelled by a normal distribution, Kolmogorov-Smirnov test was used. Comparisons between two independent groups (stage II tumors versus stage IV tumors) were performed with non-parametric Mann-Whitney U-test. Correlations with parametric variables were performed with Pearson test (P) and with non-parametric variables with Spearman test (S). Association analysis between SIRT1 protein levels and cytoplasmic β−catenin levels was performed with Chi-square test, considering median of cytoplasmic β−catenin Hscore or SIRT1 expression levels as cut-off point to separate samples between high-or low-expression levels.

Analysis between two sample groups were performed with Student’s t test, and for multiple comparisons ANOVA with Bonferroni’s post-test was used. All statistical analyses were performed with SPSS IBM software v.24. P-values<0.05 were considered statistically significant.

### Ethics

Patient samples were derived from surgical removal at Fundación Jimenez Diaz University Hospital, General and Digestive Tract Surgery Department: 95 patient samples diagnosed with stage II CRC. The Institutional Review Board (IRB) of the Fundación Jimenez Diaz Hospital, reviewed and approved the study, granting approval on December 9, 2014, act number 17/14.

Colon cancer tissues were collected using protocols approved by the corresponding Ethics Committees and following the legislation. All patients gave written informed consent for the use of their biological samples for research purposes. Fundamental ethical principles and rights promoted by Spain (LOPD 15/1999) and the European Union EU (2000/C364/01) were followed. All patients’ data were processed according to the Declaration of Helsinki (last revision 2013) and Spanish National Biomedical Research Law (14/2007, July 3).

## Acknowledgements

We thank Sergio Ruiz Reyes for excellent technical assistance.

## Author contributions

José Manuel García-Martínez, conceptualization, data curation, formal analysis, validation, investigation, methodology; Ana Chocarro-Calvo, resources, formal analysis, supervision, investigation; Javier Martínez-Useros, formal analysis, investigation, methodology; Nerea Regueira-Acebedo: methodology and formal analysis, María Jesús Fernández-Aceñero, formal analysis, validation, visualization, methodology; Alberto Muñoz, conceptualization, supervision, writing review and editing; María Jesús Larriba, supervision, review and editing; Custodia García Jiménez, conceptualization, resources, data curation, formal analysis, supervision, funding acquisition, validation, investigation, visualization, writing original draft, project administration, writing - review and editing.

## Supporting information

**Supplementary Figure S1:**
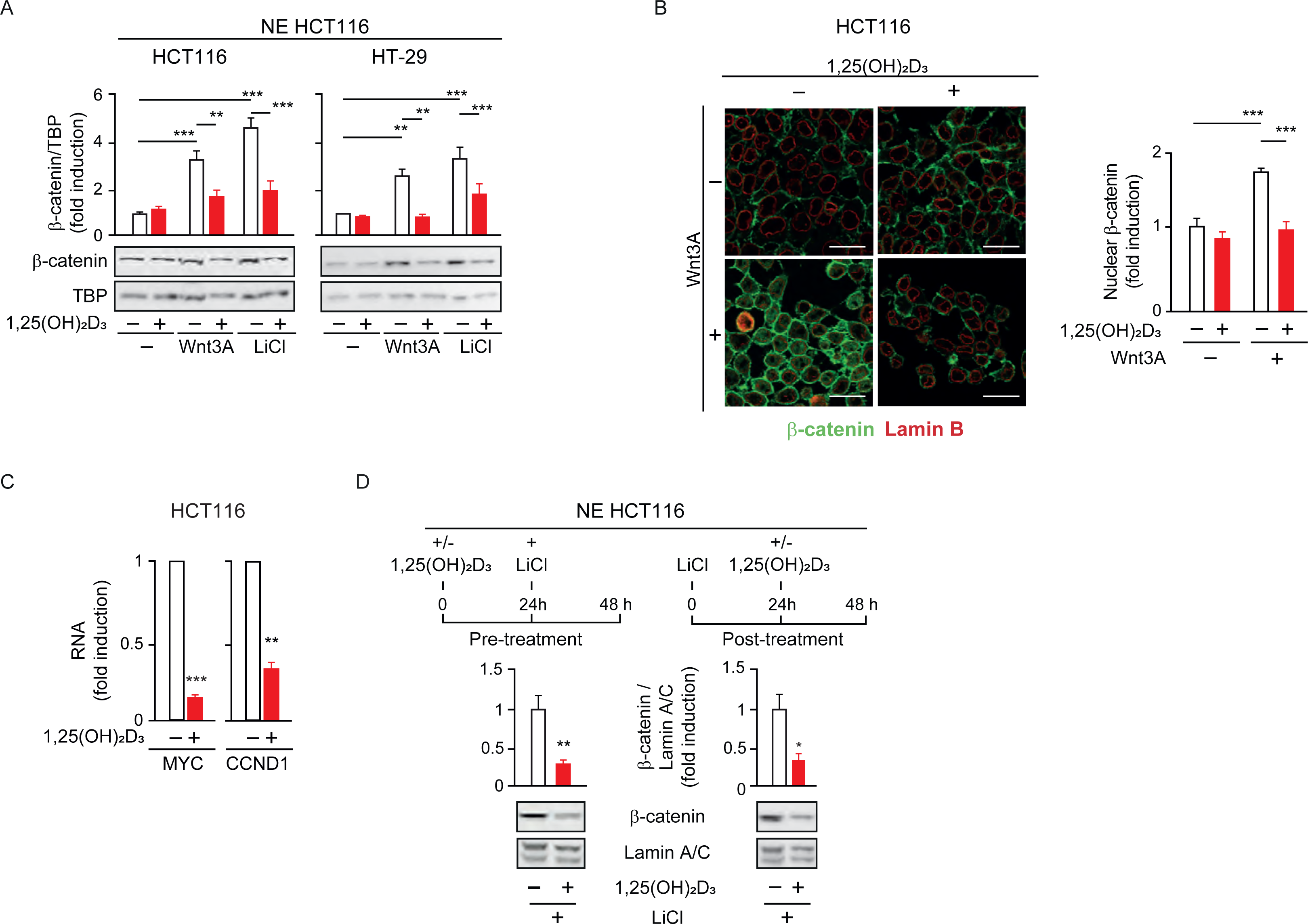
Vitamin D re locates β-catenin oppositely to Wnt and glucose and independently of whether is added before or after in CRC cells. HCT 116 or HT-29 colon cancer cells were cultured in DMEM and treated with Wnt3A (100 ng/ml) or LiCl (40nM) where indicated, before addition of 1,25(OH)2D3, 100 nM for the last 24 h. Extracts were fractionated, and nuclear extracts (NE) were analyzed. (S1A) Western-blot analysis of 1,25 (OH)2D3 effects on nuclear β-catenin levels in CRC cells treated as indicated. TBP as loading control. Representative blots and statistical analysis. (S1B) Confocal imaging of nuclear β-catenin accumulation by Wnt3A in HCT 116 cells and its reversal by 1,25(OH)2D3, representative photographies. Scale bars: 25µm. Right panel: fluorescence intensity was quantified in 3 independent experiments using ImageJ software and for each experiment, 3 different fields were evaluated per slide. (S1C) RT-qPCR of CYCLIN D1 (CCND1) and MYC from HCT 116 cells. Values normalized with endogenous control (18S) are referred as fold induction over cells without 1,25 (OH)2D3. (S1D) Order of addition analysis of the effects of 1,25 (OH)2D3 on nuclear exclusion of β-catenin in CRC cells analyzed by western-blotting. Lamin A/C were used as loading controls. Left: cells were treated with 1,25 (OH)2D3 for 24 h and then treated or not LiCl to mimic Wnt signalling. Right: cells were first treated with LiCl and then with1,25 (OH)2D3. Statistical analysis of 3 independent experiments by One-Way ANOVA in (A) or Student t-test (B) to (D); values represent mean ± SEM of triplicates; * p< 0.05; ** p< 0.01; *** p< 0.001.

**Supplementary Figure S2:**
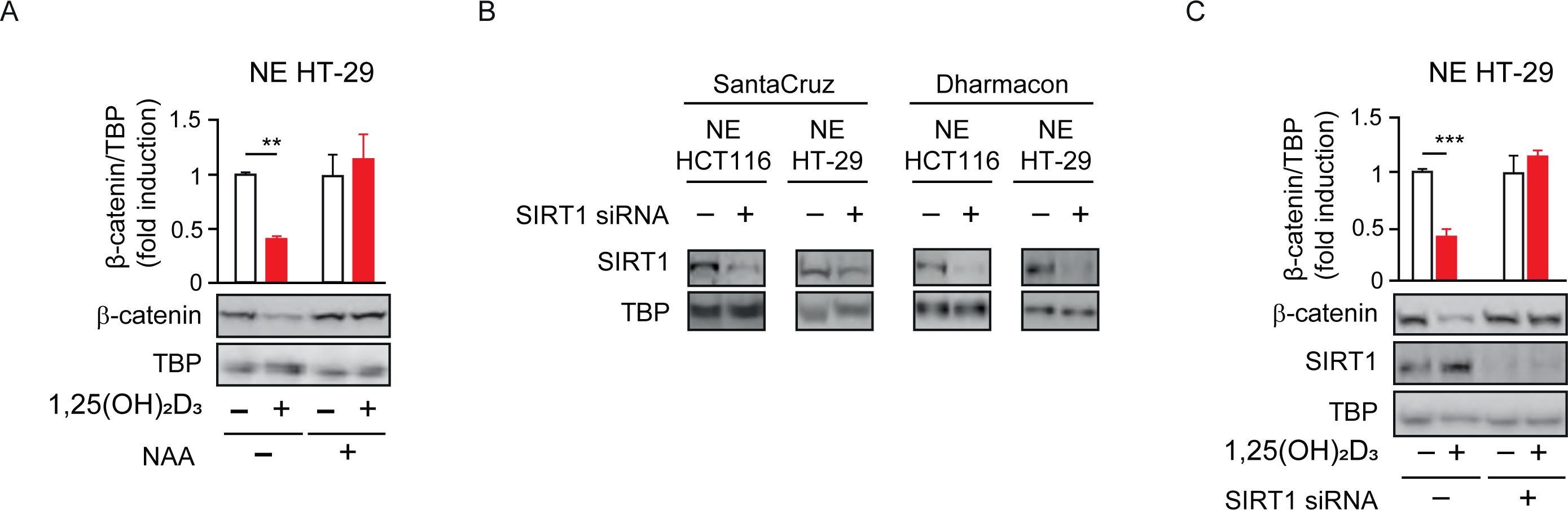
CRC cell lines respond similarly to SIRT inhibition with NAA, or SIRT1 depletion using several SIRT1 specific siRNAs. Cells were cultured and treated as in previous figures. (S2A) Western-blot analysis of the effects of 1,25 (OH)2D3 and the sirtuin inhibitor nicotinamide (NAA) on β-catenin nuclear levels; TBP as loading control. Representative blots and statistical analysis. (S2B) Comparison of SIRT1 depletion using an siRNA from Santa Cruz Biotechnology and a mixture of siRNAs from Dharmacon. Representative western blots. TBB serves as loading control. (S2C) Western-blot analysis of the effect of SIRT1 depletion by siRNAs on nuclear β-catenin levels in HT-29 CRC cells. Representative picture and statistical analysis. Statistical analysis by One-Way ANOVA of 3 independent experiments; values represent mean ± SEM of triplicates; * p< 0.05; ** p< 0.01; *** p< 0.001.

